# Time-restricted foraging under natural light/dark condition shifts the molecular clock in the honey bee, *Apis mellifera*

**DOI:** 10.1101/242289

**Authors:** Rikesh Jain, Axel Brockmann

## Abstract

Honey bees have a remarkable sense of time and individual honey bee foragers are capable to adjust their foraging activity with respect to the time of food availability. Although, there is plenty of experimental evidence that foraging behavior is guided by the circadian clock, nothing is known about the underlying cellular and molecular mechanisms. Here we present a first study exploring whether the time-restricted foraging under natural light-dark condition affects the molecular clock in honey bees. In an enclosed flight chamber (12m × 4m × 4m), food was presented either for 2 hours in the morning or 2 hours in the afternoon for several consecutive days and daily cycling of the two major clock genes, *cryptochrome2 (cry2)* and *period (per)*, were analyzed in three different tissues involved in feeding-related behaviors: brain, antennae and subesophageal ganglion (SEG). We found that morning and afternoon trained foragers showed significant phase-differences in the cycling of both clock genes in all three tissues. Furthermore, the phase-differences were more pronounced when the feeder was scented with the general plant odor linalool. Our results clearly demonstrate that foraging time functions as a strong circadian Zeitgeber in honey bees. More surprisingly our results suggest that foraging time might have the potential to override the entrainment effect of the light-dark cycle.

## INTRODUCTION

Honey bee foraging has been one of the best studied and most fruitful behavioral paradigm in the study of sensory and cognitive capabilities of insects and animals in general [1–3]. Foragers search for new food sites and continue to visit a highly rewarding food source for days and weeks till it gets exhausted. This flower constancy enables researchers to train honey bee foragers to an artificial sugar-water feeder which then can be used as an experimental tool. For example, presenting the feeder at a specific time during the day demonstrated that honey bees can be time-trained and showed for the first time that animals have a sense of time [4–6].

Since then, many behavioral studies followed investigating time-memory (= foragers are trained to visit at different feeders at different times of the day) or time-restricted foraging (= foragers are restricted to visit only one feeder presented at a specific time of the day). Time-memory experiments showed that honey bee foragers are capable of associating food related cues like odor, color or spatial location with time [7–9] and can memorize up to nine different feeder times per day [10]. Time memory and daily foraging rhythm of bees are regulated by the circadian clock [4,11].

We were interested in the phenomenon of time-restricted foraging and the question whether food-reward or foraging-related cues can function as Zeitgebers for the circadian clock in honey bees [12–14]. It is important to note that honey bee foragers do not feed but only collect the food and deliver it to the colony food stores. Previous experiments on restricted foraging in honey bees have demonstrated that foragers show food anticipatory behavior [15–17]. The accuracy of anticipation varies with time of the day; afternoon trained foragers are less accurate than morning trained foragers [15]. Furthermore, Frisch and Aschoff [18] showed that under constant light and temperature conditions in a flight room, restricted feeder presentation was capable of entraining the daily foraging activity rhythm indicating that the time of feeding has an effect on the endogenous clock. It is important to note that time-restricted feeder presentation in honey bees have clear ecological underpinnings, as there is ample evidence that flowering of many plants shows diurnal rhythmicity [19–21].

Studies on the mechanisms of food entrainment gained a lot of interest in recent years [22–24]. In vertebrates, food entrainment has a strong effect on peripheral clocks in organs like stomach, liver, adrenal glands etc. but does not affect the suprachiasmatic nucleus (SCN), the central brain clock [22,25,26]. Similarly, food entrainment experiments in *Drosophila* showed effects on peripheral clocks in fat bodies but no effect on the brain clock [27]. To test the effect of restricted food availability on the molecular circadian clock in honey bees we used the experimental protocol established by Frisch and Aschoff [18]. However, different to them we performed the studies in an outdoor flight enclosure in the presence of natural cycles of light, temperature and other environmental variables. This allowed us to test whether bees could synchronize their foraging activity to time-restricted feeder presentation while being in the presence of naturally varying environmental factors. To determine the effect of restricted foraging on the molecular clocks we monitored the daily oscillations of *cry2* and *per* mRNA expression levels for whole brains, the SEG and the antennae [28]. Unexpectedly, our study produced two challenging findings. First, the cycling of the molecular clock in the brain, SEG, and antennae of honey bees can be shifted by restricted feeder presentation under natural LD cycle. This finding indicates that food/foraging time can function as a potent Zeitgeber in honey bees. Second, odor marking of the feeder increased the entrainment effect. To our knowledge, such an effect has not been described so far. Once again, honey bees appear to have evolved intricate, and perhaps unique, clock mechanisms in the context of social organization of individual behavior.

## RESULTS

### Morning- and afternoon-trained foragers showed phase differences in the daily expression rhythm of clock genes in the brain

During ad-libitum feeding, both *cry2* and *per* expression cycles peaked around midnight as was shown before (n=5 or 6 per time point; 6 time points) (Fig 1) [28]. When colonies were restricted to 2 hour feeding regimes, either during morning or afternoon, the peaks (=acrophases) of daily *cry2* and *per* mRNA rhythm were shifted (n=4-6 per time point; 6 time points; 3 repeats) (Fig 1A and 1B). In morning-trained foragers, *cry2* mRNA level peaked shortly before midnight while in afternoon-trained, it peaked during early morning. There was on average a 5.5±1.4 hours (Watson-Williams F-test, *p* < 0.005) phase difference in *cry2* and a 7.7±4.6 hours (Watson-Williams F-test, *p* < 0.050) in *per* mRNA rhythm between morning and afternoon trained foragers (Fig 1C). In all experiments, *cry2* mRNA level oscillated with higher amplitude than *per* (S1 Fig, S1 and S2 Table). Correspondingly, daily *cry2* mRNA oscillations were significant in all the experiments while *per* was rhythmic only in 4 out of 6 experiments similar to earlier studies (S2 Table) [29].

**Fig 1.**
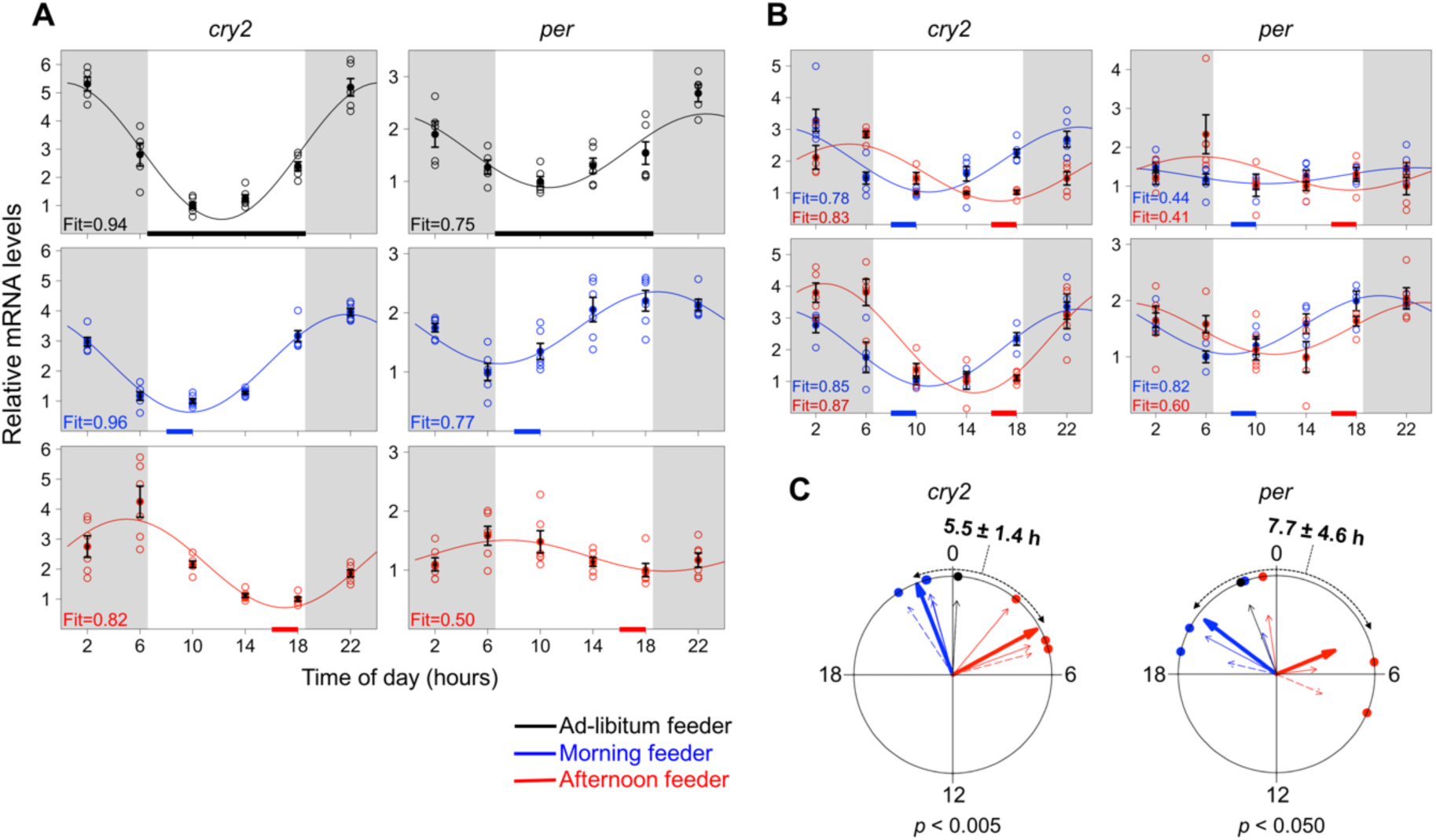
Time of feeding shifts the molecular clock in brain. (A-B) Each open circle corresponds to an individual qPCR readout from a single brain normalized with the internal reference gene, *rp49* (n=4-6 per time point). Filled circles represent the mean±SEM. Continuous line through the data points is the best fitted cosine model with fit value at the bottom left of each plot (A) Two different colonies exposed to different feeding regimes (morning or afternoon) were compared (B) Both morning and afternoon trainings were done with foragers from the same colony in subsequent experiments. (C) Acrophases (peaks) of the cosinor curves for the experiments in A and B are presented as a small dot on periphery of circular plots with a thin arrow pointing towards each dot. Units of circular plots correspond to the time in the real world, where 0 represents midnight. Acrophases from the experiment done with different colonies (shown in A) are indicated with dashed thin arrows. Lengths of the thin arrows correspond to the fit values of cosine models. Thick arrows represent the mean acrophase and length of the thick arrows correspond to coherence of all 3 acrophase values. Watson-Williams F-test was performed to compare the acrophases.

### The phases of daily clock genes expression rhythms in peripheral tissues processing food-related cues (antennae and SEG) were similarly shifted

Firstly, we analyzed the daily changes in expression of 4 major clock genes, *cry2, per, cycle (cyc)* and *clock (clk)*, in antennae and SEG of ad-libitum fed foragers (S2 Fig). Both SEG and antennae have important functions in feeding, food search and foraging [30,31]. Except *Clk*, mRNA levels of all the genes showed daily rhythmicity in both tissues (S1 Table) [32]. Later we compared these peripheral clocks in morning- and afternoon-trained foragers. The *cry2* rhythms in SEG (n=2 or 3 per time points: 6 time points; 3 repeats) were different by 4.1±0.7 hours (*p* < 0.005) (Fig 2A-2C) while in antennae (n=2-4 per time points; 6 time points; 3 repeats) this difference was 4.7±0.9 hours (*p* < 0.005) (Fig 2D-2F). The phase-differences in *per* mRNA rhythm between morning and afternoon foragers were 6.5±1.9 hours (*p* < 0.005) and 6.4±0.5 hours (*p* < 0.005) in SEG and antennae respectively.

**Fig 2.**
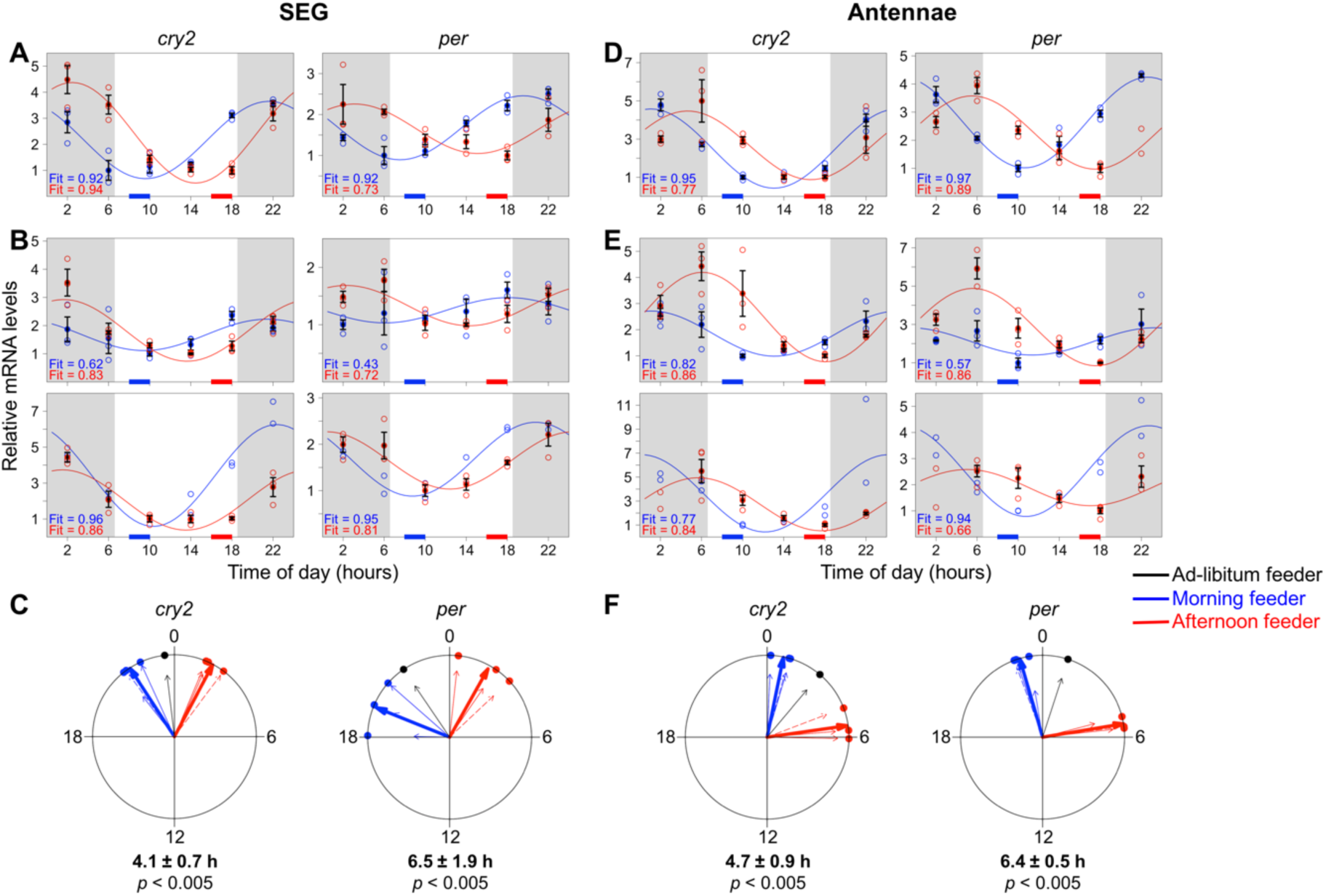
Time restricted feeding/foraging shifts the peripheral clocks in honey bee foragers. (A, B, D and E) Open circles are individual qPCR readouts, either from 4 pooled antennae or from 2 pooled SEGs normalized with the internal reference gene, *rp49* (For SEG, n=2 or 3 per time point and for antennae, n=2-4). Mean+SEM are presented with filled circles for n ≥ 3. (A and D) Different colonies were used for morning and afternoon feeding experiments. (B and E) Both morning and afternoon training were done with the same colony in subsequent experiments. (C and F) Acrophases (peaks) of the cosinor curves are presented as a small dot on periphery of circular plots with a thin arrow pointing towards each dot. Units of circular plots correspond to the time in the real world, where 0 represents midnight. Experiments, in which different colonies were used for morning and afternoon training (A and D), are depicted with dashed thin arrows. Lengths of the thin arrows correspond to the fit values of cosine models. Thick arrows represent the mean acrophase and length of the thick arrows correspond to coherence of all 3 acrophase values. Watson-Williams F-test was performed to compare the acrophases.

### Time restricted foraging to an unscented feeder is sufficient to shift the molecular clocks

The 2-hour time-restricted feeding experiments, as previously done, were performed without using any scent on the feeder. Again, on day 11, regularly visiting foragers were consecutively collected at four hours intervals (n=4-6 per time points; 6 time points; 3 repeats) from the colony and clock genes mRNA levels in the brain, antennae and SEG were measured (S3 and S4 Fig). The phase of clock genes expression rhythms in all three tissues of morning foragers were different from the afternoon foragers (Fig 3). The mean acrophases of *cry2* rhythms were 3.3±1.5 hours (*p* < 0.050) hours apart in the brain, 1.9±0.5 hours (*p* < 0.005) hours apart in the SEG and 2.3±2.7 hours (*p ≥* 0.050 or ns) hours apart in the antennae. The acrophases of *per* rhythm were different by 5.8±2.9 hours (ns) in the brain, 3.1±1.0 hours (*p* < 0.005) hours in the SEG and and 3.6±3.3 hours (ns) in the antennae. Interestingly, these phase-differences were not as pronounced as the scented feeder experiment.

**Fig 3.**
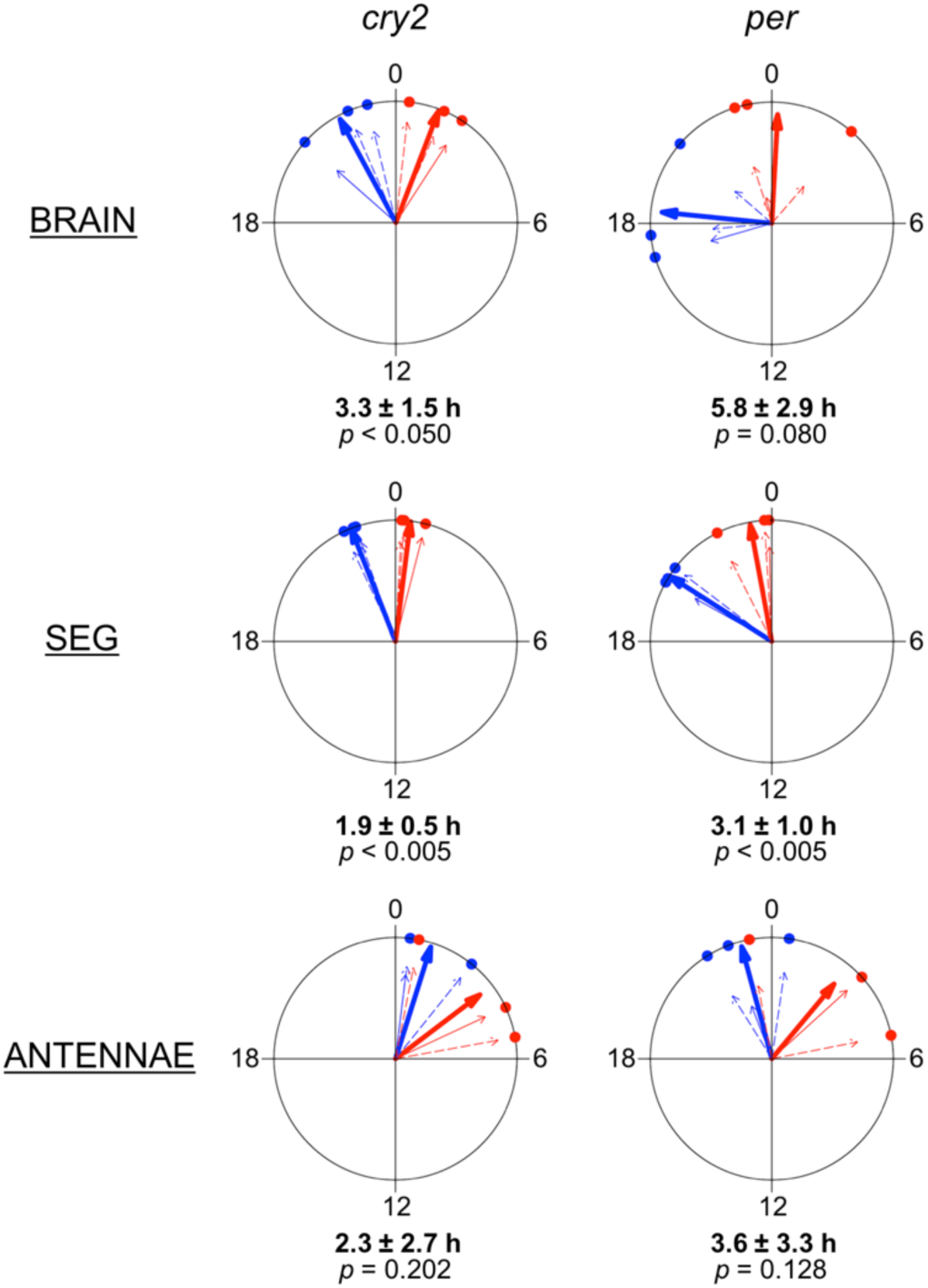
Restricted foraging at unscented feeder is sufficient to shift the molecular clock. Morning or afternoon restricted feeding was performed without odor and differences in clock genes rhythm in brain, SEG and antennae were analyzed. Only the acrophases of circadian gene expression data (S3 and S4 Fig) are presented as small dots on the periphery of circular plots with a thin arrow pointing towards each dot. Units of circular plots correspond to the time in the real world, where 0 represents midnight. Acrophases from the experiment done with different colonies are indicated with dashed thin arrows. Lengths of the thin arrows correspond to the fit values of the cosine models. Thick arrows correspond to the mean acrophase and length of the thick arrows correspond to coherence of all 3 acrophase values. Watson-Williams F-test was performed to compare the acrophases.

## DISCUSSION

Our results demonstrate that under natural LD conditions, time-restricted foraging modulates the expression of major clock genes in honey bee foragers. *cry2* and *per* mRNA expression rhythm in the brain, SEG, and antennae showed a strong phase-difference between morning and afternoon trained foragers. This finding suggests that food-reward and/or food-reward induced foraging activity are capable of shifting the molecular clock and even more importantly in the presence of the natural LD cycle. Furthermore, comparison of the clock gene cycling in foragers trained to a scented or a non-scented feeder indicates that food-reward and/or food-reward dependent foraging activity alone is sufficient to induce the changes in the molecular clock. However, the presentation of odor at the feeder led to an increase in the phase shift indicating a direct or indirect effect of the olfactory system on the molecular clock.

Our findings are supported by the results of three previous studies. Thirty years ago, Frisch and Aschoff [18] demonstrated that restricted feeding under constant light and temperature conditions in a flight room entrains the daily foraging activity of honey bee colonies. More recently, Naeger et al. [33] showed that foragers of the same colony visiting a feeder either in the morning or in the afternoon differ in the expression levels of *cry2* and *per*. These differences nicely correlate with our data on the phase-shift with morning and afternoon foraging. Finally, Krauss et al. [20] provided evidence that honey bee foragers have preferred activity times, similar to human chronotypes [34], and this “shift work” among foragers has a genetic basis.

In contrast to honey bees, *Drosophila* flies apparently do not show any signs of food entrainment under LD cycle [35]. Under constant dark conditions restricted feeding entrained the molecular clock in the fat bodies but not in the brain [27]. In mammals, restricted feeding under LD conditions entrained the molecular clock in several brain regions as well as in various peripheral organs, e.g. liver, stomach, adrenal glands but not the central clock in the SCN [22,26]. Our whole brain qPCR studies showed that restricted feeding affects clock gene expression in the brain. Thus, we cannot exclude the possibility that these changes occurred in brain cells, e.g. glia, that do not strictly belong to the neuron populations representing the central insect brain clock [36]. On the other hand, studies in Drosophila and honey bees suggest that behavioral activity rhythms are regulated by the central brain clock [36–39].

In our experiments, directions of the phase-shift induced by morning or afternoon restricted foraging were similar for all three tissues tested. Morning restricted foraging shifted the peak to an earlier time compared to unrestricted foraging whereas afternoon restricted foraging shifted the peak to a later time. Our experiments do not allow any conclusion about whether the clocks in the three tissues are synchronized (S5 Fig) which each other or whether they are independently affected by food cues or foraging activity.

Although there are ample evidences for food entrainment, there haven’t been any reports, to our knowledge, on the effect of food-related odors on endogenous clock. In our experiments, we observed that the presence of feeder associated odor results in stronger phase-shift (S6 Fig). Odors play a key role in the recruitment of honey bee foragers to food sites and the identification of a learned food source [6,19,40–42]. Moreover, honey bees are capable of associating odors with a time of the day [10]. While the underlying mechanism of honey bee time-memory is not well understood yet, we propose that the floral odor may play an important role in time-learning by influencing the endogenous clock. The neuronal pathway/s through which the olfactory signals interconnect with the molecular clocks in the brain is yet to be discovered.

Given what is known about food entrainment in vertebrates and flies, our findings suggest that honey bee are different or even special with respect to the effect of food or foraging on the circadian clock. If so, why are they different? Likely, most solitary invertebrates and vertebrates adjust the major activity rhythms according to the LD cycle being active during the day or the night. Food search is only one part of that activity and food availability is mostly not very predictable. In case it is predictable, anticipatory physiological and behavioral processes can be regulated by different oscillators, whereas the main clock follows the LD cycle. In contrast, honey bee foraging is a social behavior with the search for food and foraging being highly socially organized and energetically optimized [43]. Different tasks of this process are performed by specialized worker groups (e.g. scouts, recruits, receivers, food storers), and each worker group mainly does only one task during the whole day. More impressively, foragers even specialize on a specific food patch or food plant (flower constancy [44]). During the flowering seasons, food is generally available in plenty during the whole day. Given that honey bee colonies have hundreds to thousands of foragers, it could be an energetically optimized strategy to send out foragers for only part of the time, like shift workers in human societies. To do so, honey bees might have evolved two mechanisms – 1) they might have evolved different chronotypes with preferences of being active in the morning or in the afternoon, and 2) they might have evolved the capability to entrain their activity rhythm with respect to the flowering time of the learned food plant.

**Fig 4.**
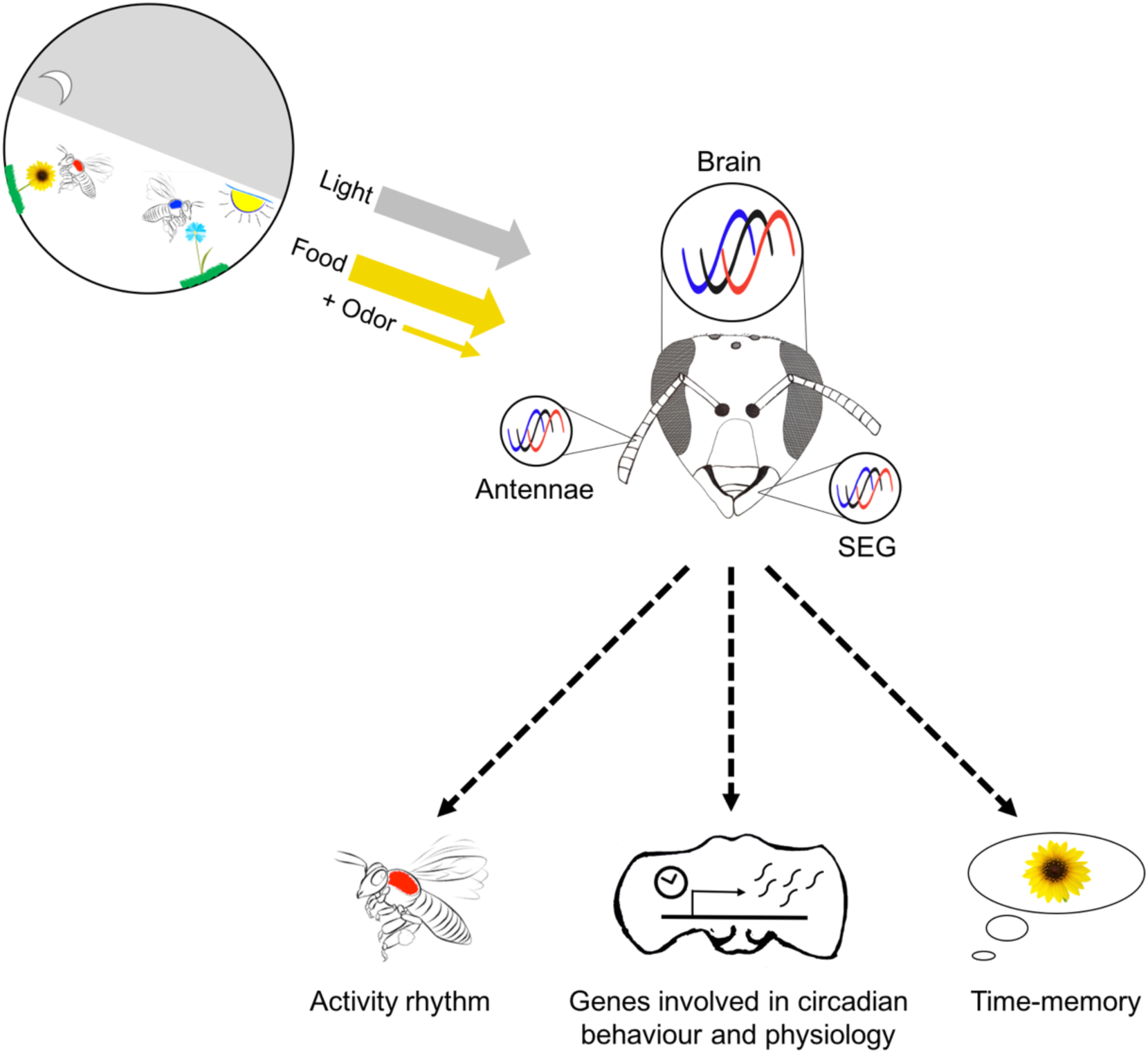
Schematic summary outlining the proposed mechanism of food anticipation in honey bees. In honey bees, food-reward or food-reward dependent foraging behavior can function as a Zeitgeber for molecular clocks in the brain, SEG, and antennae. Furthermore, this food/foraging Zeitgeber appears to have potential to over-riding the light/dark Zeitgeber. Addition of odors to the food increased the effect of the food entrainment. We propose that food/foraging entrainment affects (1.) daily activity rhythm [18,45], (2.) expression pattern of clock controlled genes [33,46], some of them involved in foraging associated processes (e.g. olfactory and visual sensitivity [33,47]), and (3.) time-memory [33].

## MATERIALS AND METHODS

### Animals

*Apis mellifera* colonies with a naturally mated queen obtained from local beekeepers and maintained on the campus of the National Centre for Biological Sciences (NCBS-TIFR), Bangalore were used for all experiments. For the time training experiments, colonies were kept in an outdoor flight enclosure (12m × 4m × 4m), which allowed regulated presentation of food sources such as a sugar-water feeder and separate pollen feeder, under naturally fluctuating environmental conditions. Colonies were allowed to adjust to the flight cage for one week before beginning the time restricted feeder presentation. During this week the sugar and pollen feeder was available ad-libitum.

### Time restricted feeder presentation

Time-restricted feeder training was carried out by presenting sucrose feeder (1M sucrose solution) for 2 hours either in the morning (8.00 to 10.00) or the afternoon (16.00 to 18.00) for 10 consecutive days. Restricted feeding experiments were performed with scented as well as unscented feeders. For scented feeders, 5ul drop of 100 times diluted (in mineral oil) linalool racemic mix (Sigma-Aldrich) were kept on 4 filter papers (70mm diameter) equally distant from the sucrose feeder plate. Feeder was washed and scented filter papers were replaced with fresh filter papers after the training time every day. On the 7th and 8th day, 4 and 3 days before the collection, foragers visiting the feeder were marked with two different colors to ensure that we will collect foragers that were frequently visiting the feeder. On the 11th day, without presenting the feeder, marked foragers were serially collected at 6 different time points i.e. 6.00, 10.00, 14.00, 18.00, 22.00 and 2.00 from the colony. Night collections were done using dim red light. Collected bees were flash frozen in liquid nitrogen and stored in a −80°C deep freezer.

### Dissection, RNA isolation and qPCR

Antennae, brains and SEGs were dissected on dry ice. During brain dissections, ocelli, compound eyes and all glandular tissue were removed. Total RNA from antennae (4 antennae/sample), SEGs (2 SEGs/sample) and brains (1 brain/sample) was isolated using Trizol^®^ reagent (Invitrogen) following the manufacturer’s protocol. RNA quantity and quality were assessed via nanodrop measurements and agarose gels. All samples were treated with DNAse (DNase I, Amplification Grade, Invitrogen). 0.5ug-1ug of RNA was used to generate cDNA using Superscript III (Invitrogen) and oligo d(T)16 primers (Invitrogen). Primer pairs (S3 Table) for qPCR were designed in a way that one of the primers covered exon-exon junctions, not allowing the amplification of genomic DNA if any. Each reaction was run in triplicates using (10ul reaction mix) KAPA SYBR FAST qPCR Master Mix (Sigma-Aldrich) in Applied Biosystems 7900HT Fast Real-Time PCR system. Purity of all the qPCRs was verified using dissociation/melt curve analysis. Although all primers showed efficiency of 95–100%, standard curves with a separate stock cDNA were generated in each qPCR (384-well) plate to reduce inter run variability. Test genes and *ribosomal protein 49 (rp49)* levels in all 6 time point samples from the same experiment were analyzed in the same plate. mRNA levels quantification was based on the linear values interpolated from the standard curves. *rp49* used as internal control gene for normalization.

### Statistical analysis

R [48] was used for all the statistical analysis. 24-hour cosinor models were incorporated in gene expression data and acrophase and fit values were calculated using cosinor package [49]. In addition, non-parametric JTK cycle analysis [32] was also performed to detect significance and amplitude of 24-hour mRNA oscillations (Table S1 and S2). Watson-Williams F-test was performed to analyze statistical differences in acrophase values.

## AUTHORS’ CONTRBUTIONS

R.J. and A.B. designed the experiments. R.J. performed the experiments and analyzed the data. R.J. and A.B. wrote the manuscript.

## ACKNOWLEDGEMENTS

We thank Ravi Boyapati for helping in sample collections. We also thank Sheeba Vasu, Sruthi Unnikrishnan and members of the bee group for valuable comments on earlier versions of the manuscript. R.J. was supported by Indian Council of Medical Research (ICMR) fellowship; A.B. is supported by NCBS-TIFR institutional funds No. 12P4167.

